# The mechanism of sphingolipid processing revealed by a GALC-SapA complex structure

**DOI:** 10.1101/112029

**Authors:** Chris H. Hill, Georgia M. Cook, Samantha J. Spratley, Stephen C. Graham, Janet E. Deane

## Abstract

Sphingolipids are essential components of cellular membranes and defects in their synthesis or degradation cause severe human diseases. The efficient degradation of sphingolipids in the lysosome requires lipid-binding saposin proteins and hydrolytic enzymes. The glycosphingolipid galactocerebroside is the primary lipid component of the myelin sheath and is degraded by the hydrolase β-galactocerebrosidase (GALC). This enzyme requires the saposin SapA for lipid processing and defects in either of these proteins causes a severe neurodegenerative disorder, Krabbe disease. Here we present the structure of a glycosphingolipid-processing complex, revealing how SapA and GALC form a heterotetramer with an open channel connecting the enzyme active site to the SapA hydrophobic cavity. This structure defines how a soluble hydrolase can cleave the polar glycosyl headgroups of these essential lipids from their hydrophobic ceramide tails. Furthermore, the molecular details of this interaction reveal how specificity of saposin binding to hydrolases is encoded.

## Introduction

Sphingolipids are both essential membrane components and bioactive metabolites that regulate critical cell functions. Defects in sphingolipid metabolism underlie a range of diseases, including lysosomal storage diseases, and are implicated in a number of cancers (1–3). Sphingolipid degradation occurs in the lysosome and depends upon two families of proteins: glycosyl hydrolases, and lipid-transfer proteins including saposins and GM2 activator proteins (4–7). The hydrolases are water soluble while the substrates are embedded in lysosomal intraluminal vesicle membranes. The steric crowding of headgroups and lateral association of sphingolipids into clusters prevents hydrolases from accessing the scissile bonds of their target substrates. Saposins are required to facilitate sphingolipid processing by hydrolases, and extensive work provides evidence for two general models of saposin action (8–11). The ‘solubiliser’ model proposes the complete extraction of lipids from the membrane, while the ‘liftase’ model envisages saposin proteins binding directly to compatible membranes and improving lipid accessibility by membrane distortion, destabilisation or localised remodelling. The saposin proteins are produced as a polycistronic prosaposin protein that, upon delivery to the lysosome, is cleaved into the four saposins: A, B, C and D (12–14). The functions of these four saposins are distinct, as they cannot compensate for the loss of each other, and saposins appear to function specifically with their associated hydrolase (15, 16). Lysosomal storage diseases, and more specifically sphingolipidoses, are caused by mutations that inhibit degradation of sphingolipids. For example, Gaucher and Krabbe diseases are caused by loss of glucocerebrosidase (GluC) and galactocerebrosidase (GALC) activity, respectively, or by mutations in their associated saposins SapC and SapA, respectively (17–20).

Several high-resolution structures of saposins have been determined to date, revealing huge conformational variability and a propensity to form oligomeric assemblies (21–25). Saposins can broadly be described as existing in two states: a “closed” monomeric form where the helical protein folds back on itself, burying a large hydrophobic core; or a more “open” dimeric form, possessing a hydrophobic cavity into which lipids and detergents can bind. A recent structure of SapA revealed that this open conformation can form lipo-protein discs (21) and these discs have recently been exploited to aid the determination of challenging membrane protein structures by crystallography and CryoEM (26). However, it has remained unclear what form of saposin-lipid complex mediates binding to their cognate hydrolase. To remedy this deficit we have solved the structure of GALC in complex with SapA, defining how saposins solubilise lipids for processing by soluble hydrolases.

## Results

Murine GALC and SapA were expressed and purified from mammalian cells and *E.coli,* respectively. Pulldown assays reveal that SapA does not bind GALC at neutral pH, equivalent to that of the endoplasmic reticulum, but instead these proteins form a complex at low pH equivalent to that of the endolysosomal compartments where lipid degradation occurs (Fig. 1A). This interaction depends upon the presence of detergents, suggesting that the “open” (dimeric) lipid-bound form of SapA mediates the interaction (Fig. 1B). The pH-dependency of the interaction, combined with the low pI of SapA (pH 4.5) and the higher pI of GALC (pH 6.1), suggests that the interaction could be mediated by electrostatic interactions. In support of this, the interaction is sensitive to high salt concentrations as the presence of increasing concentration of NaCl reduces binding in these pulldown assays (Fig. 1B).

**Figure 1.**
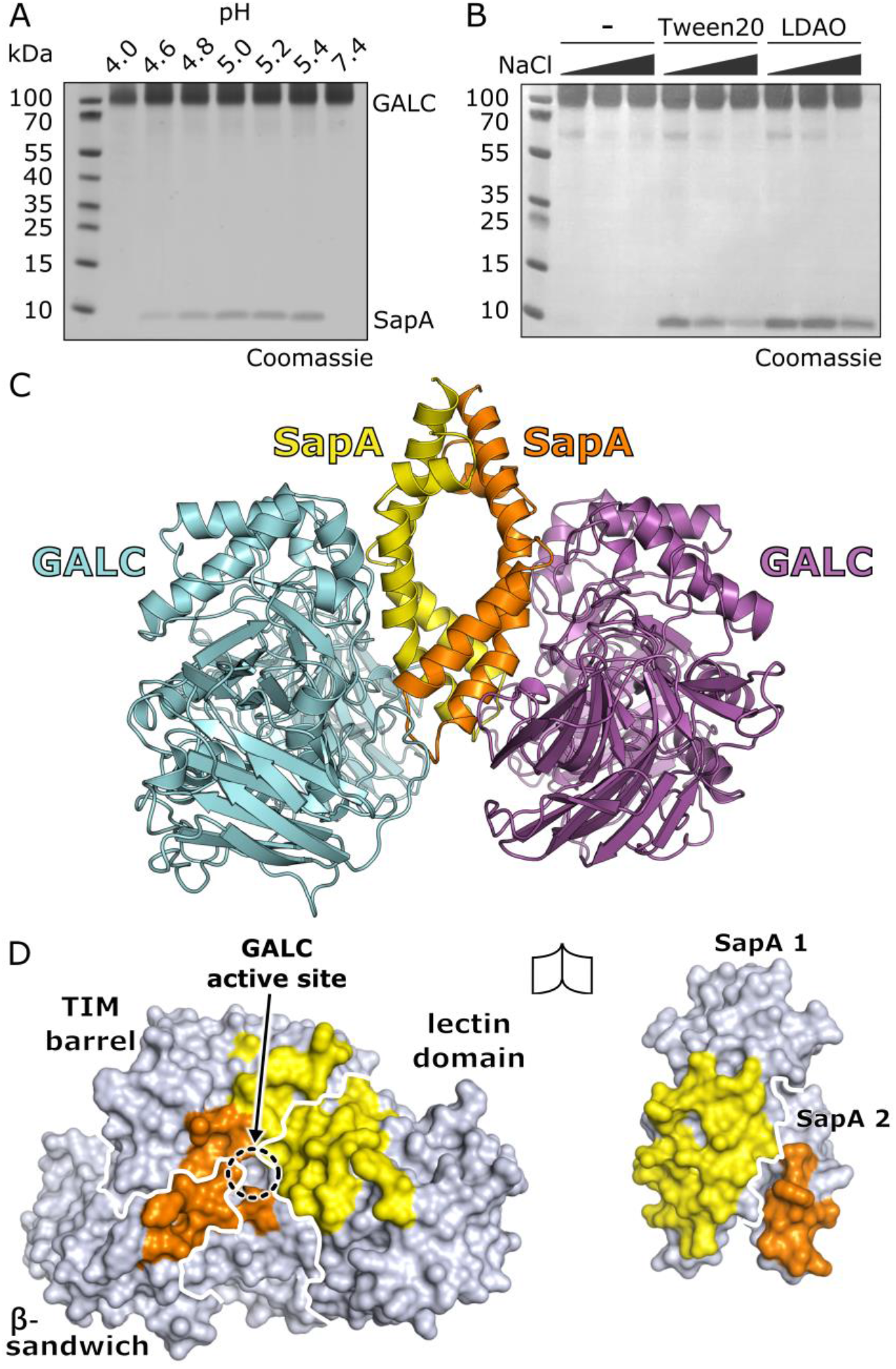
GALC-SapA complex formation and structure. (A) Coomassie-stained SDS-PAGE of pulldown assays with immobilised GALC in the presence of the detergent, Tween-20, shows binding to SapA at pH values equivalent to the late-endosomal/lysosomal compartment. Glycosylation of GALC results in a smeared band on gradient gels. (B) Pulldown assays at pH 5.6 identify that the interaction requires the presence of detergents, such as Tween 20 or LDAO, and is reduced with increasing NaCl concentration (20, 150 and 300 mM NaCl). (C) The crystal structure of the GALC-SapA complex reveals a symmetric 2:2 heterotetramer of SapA (yellow and orange) with GALC (cyan and magenta). (D) The interacting surfaces of GALC (left) and the SapA dimer (right), rotated by 90° with respect to (C) to highlight the respective interaction surfaces. On the GALC surface the interacting regions contacted by SapA chain 1 (yellow) and SapA chain 2 (orange) are highlighted, the different domains of GALC are outlined (white lines) and the active site is identified. On the SapA surface the residues that interact with GALC are highlighted for each chain and the different chains of SapA are outlined (white line).

Based on these insights, the GALC-SapA complex was formed at pH 5.0 in the presence of the detergent LDAO and the X-ray crystal structure of this complex solved and refined to 3.6 Å resolution (Fig. 1C and Table 1). The structure reveals that the complex forms a 2:2 heterotetramer, comprised of a central SapA dimer with two molecules of GALC binding symmetrically on the surface of the SapA dimer. By peeling apart the complex structure the details of the surfaces involved in the interaction between GALC and SapA are revealed (Fig. 1D). Importantly, the active site of GALC is buried in the centre of the interaction interface. Not only does the SapA dimer sit over the active site, it makes contact with all three domains of GALC: the TIM barrel, β-sandwich and lectin domains (27). This extensive interface buries a total surface area of 1366 Å^2^, of which 70% is contributed by one SapA chain and 30% by the other (yellow and orange, respectively, in Fig. 1D).

**Table 1.** Data Collection and Refinement Statistics. Values in parentheses are for highest-resolution shell.

|  |  |
| --- | --- |
| **Data collection** |  |
| Space group | *P* 6<sub>2</sub> 2 2 |
| Cell dimensions |  |
| *a, b, c* (Å) | 187.2, 187.2, 360.2 |
| $\alpha, \beta, \gamma$ (°) | 90, 90, 120 |
| Resolution (Å) | 180.12 – 3.60 (3.70 - 3.60) |
| *R*<sub>merge</sub> | 0.386 (2.946) |
| CC<sub>1/2</sub> | 0.945 (0.677) |
| *I*/σ*I* | 9.3 (1.5) |
| Completeness (%) | 100 (100) |
| Redundancy | 25.6 (25.2) |
| **Refinement** |  |
| Resolution (Å) | 162.11-3.60 |
| No. reflections | 43,825 |
| *R*<sub>work</sub>/*R*<sub>free</sub> | 0.213/0.226 |
| No. atoms |  |
| Protein | 11,522 |
| Other<sup>1</sup> | 252 |
| Water | 20 |
| *B*-factors |  |
| Protein | 100.9 |
| Other<sup>1</sup> | 139.3 |
| Water | 65.6 |
| Ramachandran |  |
| Favoured (%) | 95.7 |
| Outliers (%) | 0.5 |
| r.m.s. deviations |  |
| Bond lengths (Å) | 0.008 |
| Bond angles (°) | 0.94 |

<sup>1</sup> Includes carbohydrate, ions and LDAO molecules

Analysis of the SapA dimer identifies the presence of disordered detergent within the core. This is supported by three observations: the average electron density in the cavity is lower than the surrounding solvent, consistent with the presence of detergent (Fig. S1A); an ordered LDAO headgroup can be modelled at a point of contact with a SapA surface residue (Fig. S1B); and multiple exposed hydrophobic residues line the core, consistent with this surface being exposed to non-polar solvent (Fig. 2A, *right panel).* Further analysis of the hydrophobic surface of the SapA dimer reveals a clear hydrophobic patch at the point where the two SapA chains contact the GALC surface (Fig. 2A, *central panel,* and Fig. 1D). A cross-section through the structure at this position shows that this hydrophobic patch encircles an opening in the SapA dimer surface that lies directly opposite the GALC active site (Fig. 2B). This reveals a continuous open channel that stretches from the GALC active site through to the hydrophobic cavity buried in the SapA dimer. The hydrophobic acyl chains of glycosphingolipids such as galactocerebroside (GalCer) can thus be shielded from the polar solvent by the SapA dimer while the hydrophilic glycosyl head groups are presented to the GALC active site. Previous work from our group identified how *bona fide* substrates bind GALC and the position of this substrate is illustrated in the active site of the GALC-SapA complex (Fig. 2B). Substrate binding results in small but significant conformational changes of key residues at the GALC active site (28). Although the plasticity of the active site and the conformational flexibility of the GalCer acyl chains hindered accurate modelling of a GalCer substrate into the channel of the GALC-SapA complex, it is clear that the distance spanned by GalCer matches that of the channel dimensions (Fig. 2C). We therefore propose that the GALC-SapA structure presented here represents the functional complex that facilitates glycosphingolipid catabolism in vivo.

**Figure 2.**
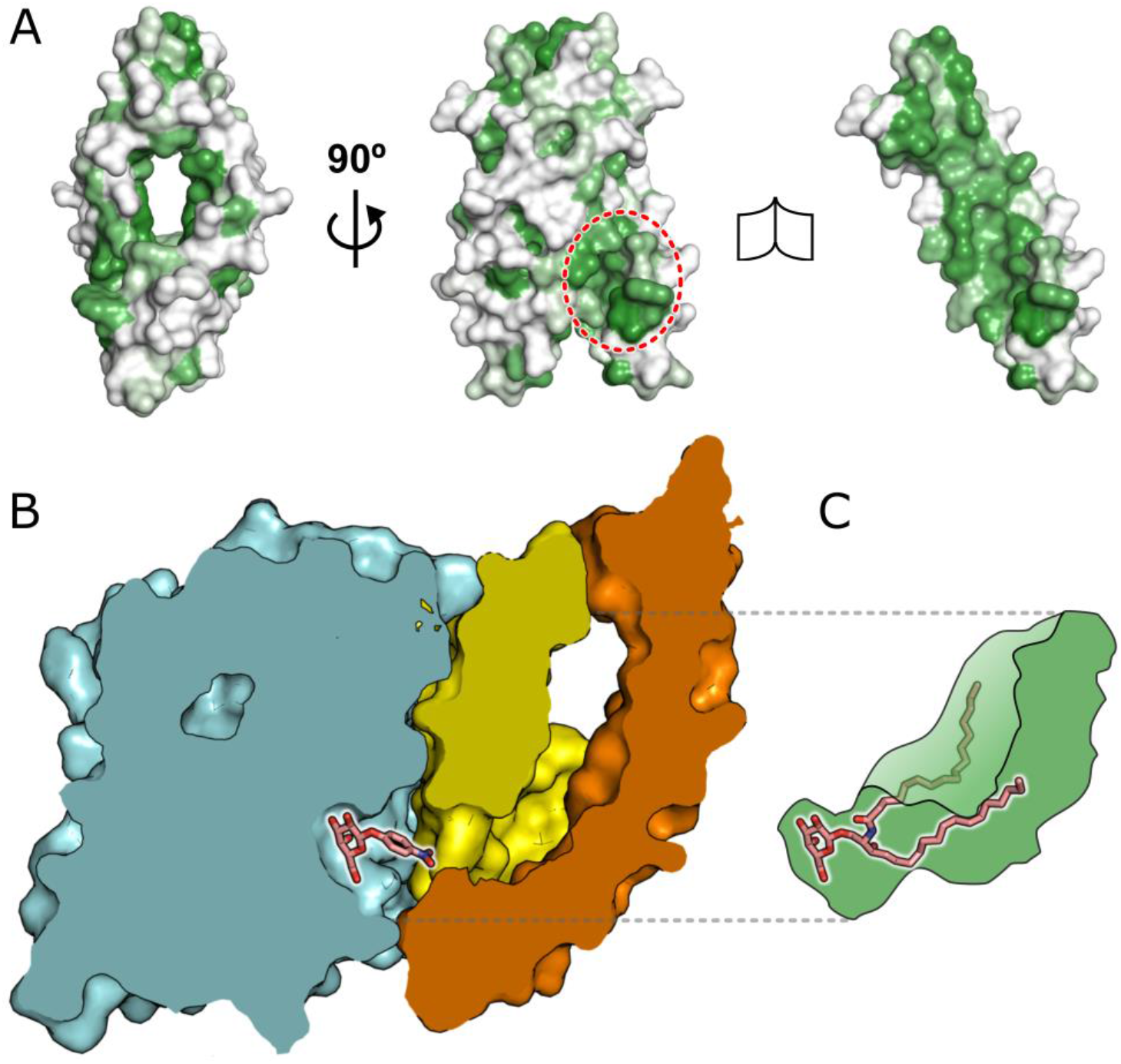
The GALC-SapA structure reveals a hydrophobic channel buried in the core of the complex. (A) Residue hydrophobicity (green) is mapped onto the surface of the SapA dimer. The left panel is in the same orientation as shown in panel 1C, while the central panel is rotated by 90° to be in the same orientation as Fig. 1D. The region that binds over the GALC active site is circled (red dashed line). The right panel maintains the orientation of the central panel, with one SapA chain removed to reveal the highly hydrophobic inner surface of the SapA dimer. (B) Cross-section through the GALC-SapA structure reveals an open channel stretching from the GALC active site into the SapA hydrophobic cavity. The GALC surface and cut-through is shown in cyan with substrate (pink sticks) positioned in the active site based on the previous structure of GALC in complex with a soluble substrate (PDB ID 4CCC (28)). The SapA dimer surface is shown in yellow and orange. For clarity, the second GALC monomer is not shown. (C) Illustration of the hydrophobic cavity (green) showing that the lipidated substrate galactocerebroside (pink sticks) fits into this cavity, bridging the channel from the GALC active site to hydrophobic SapA core.

Analysis of the molecular details at the interface reveals that several sidechains of residues along the length of one SapA chain form critical hydrogen bonds and electrostatic interactions with residues in GALC (Fig. 3A and Fig. S2). These interactions span two stretches of this SapA chain: residues 19–32 and 49–63. The interactions contributed by the second SapA chain encompass residues 36–40 and are primarily backbone hydrogen bonds, indicating that this surface is unlikely to define the binding specificity. To confirm that the interface observed in our crystal structure mediates the interaction in solution, a series of point mutations were made in SapA and tested for their ability to bind GALC (Fig. 3B, 3C). Single charge inversions at residues K19, E25, E49, D52 or D60 each abolish binding to GALC, highlighting the electrostatic nature of the interface. Inverting the charge at residue 32 significantly reduces binding but does not completely abolish binding. To ensure that alteration of surface charge alone is not sufficient to abolish binding we mutated residue E65, which does not lie in the interface, to lysine and confirmed it does not interfere with SapA binding to GALC. In addition to electrostatic interactions, hydrophobic interactions at the interface are also critical for binding. The sidechain of L28 is buried in a hydrophobic pocket of the GALC surface (Fig. S2C). Mutation of this residue to arginine completely knocks out binding to GALC. Residue N21 lies at the interaction interface and forms a backbone hydrogen bond with G144 and a number of hydrophobic interactions with the GALC surface. However, in the context of the cell this highly-conserved residue is post-translationally gylcosylated. N-linked glycans often interact with hydrophobic patches on the surface of proteins (29) and mutation of N21 to the bulky hydrophobic sidechain tyrosine moderately increased SapA binding to GALC, suggesting that the glycosylation of N21 contributes to SapA-GALC binding *in vivo.* Residue 24 is a glutamine and forms hydrogen bonds with residues T523, R520 and A525 of GALC (Fig. 3A and S2). In human SapA the corresponding residue is a glutamate, which could form a salt bridge with R520. Mutation of Q24 to lysine blocks SapA binding to GALC, supporting its role in stabilisation of the interaction. Although the only hydrogen bond formed by K19 is via the backbone to GALC residue S146, the acyl region of the K19 sidechain forms hydrophobic interactions with the GALC surface. Disruption of this hydrophobic interaction by substitution of K19 for glutamate abolishes the GALC-SapA interaction. Interestingly, in human GALC, there is a glutamate residue at position 146 rather than a serine, raising the possibility that SapA K19 could form a salt bridge with residue 146 in the human GALC-SapA complex. Taken together, our mutational analysis confirms the critical importance of SapA residues K19, Q24, E25, L28, E32, E49, D52 and D60 for complex formation.

**Figure 3.**
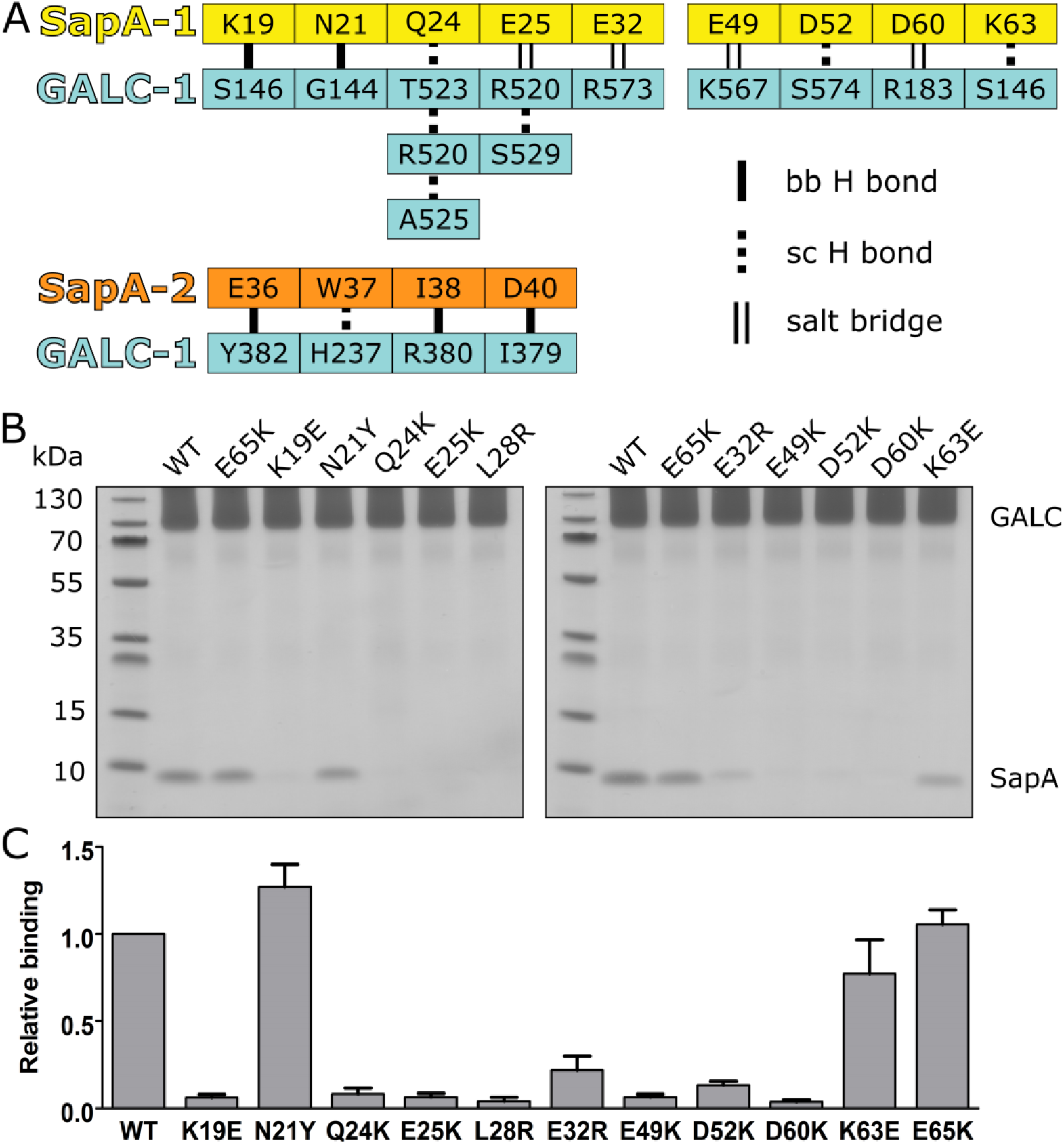
Critical residues at the GALC-SapA interface. (A) Schematic diagram illustrating the backbone (bb) hydrogen bonds, sidechain (sc) hydrogen bonds and salt bridges observed between GALC (cyan) and specific residues of SapA chains 1 (yellow) or 2 (orange). (B) Coomassie-stained SDS-PAGE of pulldown assays with immobilised GALC identifies critical SapA mutations at the interface that abolish GALC binding in solution. (C) Quantification of SapA binding in pulldown assays (n=3, error bars represent SEM).

There is high structural similarity between the different saposin proteins (30), suggesting that the specific binding to their cognate enzyme is sequence-mediated. The identification here of specific residues essential for SapA binding to GALC provides a framework for understanding this specificity. Single charge inversions at specific residues are sufficient to abolish binding *in vitro* and thus highlight potentially critical specificity determinants. Sequence alignments of saposins A-D reveal charge inversions, equivalent to those used in this study, in the region spanning residues 19–32 (Fig. 4A and 4B). Based on our binding data, any one of these amino acid changes in saposins B, C and D would block an interaction with GALC. While we were unable purify SapC, we observed that GALC binds specifically to SapA but not to SapB or SapD (Fig. 4C). This is consistent with the specificity for GALC being conferred by SapA residues 19–32, one of three stretches of SapA residues that mediate the interaction with GALC. As stated previously, the central region spanning SapA residues 36–40 primarily participates in backbone interactions so does not determine specificity. Although residues E49 and D60 of the third interacting region of SapA both form critical salt bridges at the GALC interface, the absence of charge inversions in the other saposin proteins suggests that these interactions contribute to affinity of hydrolase binding rather than specificity.

**Figure 4.**
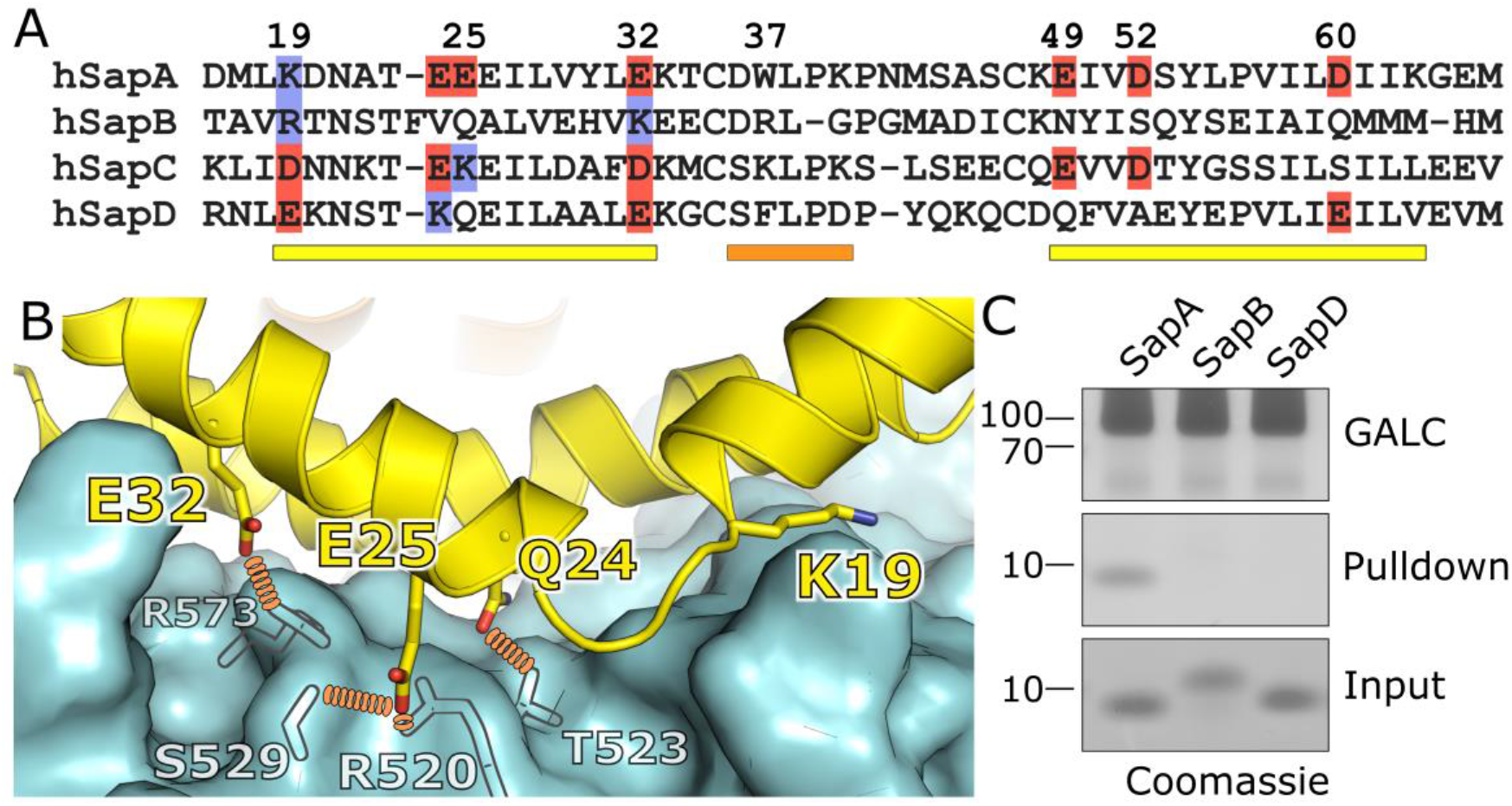
Saposin specificity. (A) Sequence alignment of human saposins A-D highlighting charge inversions of critical interface residues. Acidic residues (red) and basic residues (blue) at critical positions are highlighted. Coloured bars below the sequence identify the regions that interact with GALC: two regions from SapA chain 1 (yellow) and the one from SapA chain 2 (orange) as detailed Fig. 3A. (B) Critical residues that may determine binding specificity are shown (sticks) on SapA (yellow) at the interface with GALC (cyan surface, key residues depicted as transparent sticks). Hydrogen bonds between GALC and SapA residues are highlighted (orange dotted lines). (C) Coomassie-stained SD PAGE pulldowns with immobilised GALC illustrating specific binding of SapA, but not SapB or SapD.

## Discussion

Here we reveal molecular details of how glycosphingolipids can be bound by SapA and presented to the active site of GALC for catabolic processing. The interaction of SapA with GALC is pH dependent, consistent with its activity in the late endosomal/lysosomal compartments. The interaction also requires the presence of detergents. Structures of SapA in the absence of detergents remain closed (23, 31) whereas in the presence of detergents SapA adopts an open conformation (this study and (21)), suggesting that detergent mediates the formation of open SapA dimers essential for the interaction. The crystal structure of GALC-SapA determined here reveals the complex to be a heterotetramer with an extensive interaction interface that involves residues from all domains of GALC and both chains of the SapA dimer. There is an open channel running from the GALC active site through to the hydrophobic cavity of the SapA dimer, suggesting a clear mechanism for glycosphingolipid binding and processing.

The SapA dimer structure determined here differs from the previous structure of a SapA lipo-protein disc in isolation (21). Although the conformation of the SapA monomer is essentially identical, in our structure there are direct protein-protein contacts between the saposin monomers, whereas the previous structure contains no SapA-SapA contacts and the assembly is maintained via interactions between detergent molecules. Another difference between these SapA dimer structures is the relative orientations of the monomers. In our structure the monomers are head-to-head, with all termini together at one end, while in the lipo-protein disc structure the monomers adopt a head-to-tail orientation. Comparison of these different SapA dimers shows that the GALC-interacting surface identified in our structure is surface exposed in the lipo-protein disc structure. Docking of GALC onto the surface of the lipo-protein disc structure reveals this SapA dimer to be compatible with GALC binding (Fig. S3A). In this alternative arrangement the opening on the surface of the SapA dimer is less hydrophobic and, due to the loss of interactions with the second SapA chain, the substrate channel is more open and exposed to surrounding solvent (Fig. S3B and C). Although this alternative arrangement may be compatible with substrate binding, the tighter SapA dimer and enclosed hydrophobic channel identified in our complex provides a more protected environment from the hydrophilic solvent. We therefore conclude that our structure, possessing a large buried surface area, an enclosed hydrophobic channel and an obvious mechanism for substrate presentation to GALC, represents the *bona fide* interaction that stimulates SapA-mediated lipid catabolism in the lysosome.

The combination of structure analysis, pulldown assays and sequence comparison of the saposin family members identifies residues 19, 24, 25, 28, 32, 49, 52 and 60 of SapA as essential for binding to GALC. Within this subset, residues 19, 24, 25 and 32 are likely to be critical for determining binding specificity. The importance of the specific charges at these positions is highlighted by the loss of binding upon charge inversion and the conservation of these charges across species (30). The charge inversions between saposin proteins at these specific positions strongly suggests that the relevant patches on their cognate enzymes will possess the complementary charged residues. Despite the lack of structural similarity between glycosyl hydrolases, this identification and the observation that the saposins must sit over the active site provides the first framework for identifying the equivalent interaction interfaces on other saposin-dependent hydrolases.

While ours is the first saposin:enzyme structure, we can compare it to the recent structures of acid sphingomyelinase (ASM), which possesses an intra-molecular saposin-like domain (32–34). Although the enzymatic portion of ASM is not structurally similar to GALC the saposin-like region adopts a similar conformation to SapA in the GALC-SapA complex, allowing a reliable overlay to be made based on this portion (Fig. S4A). This comparison reveals that the catalytic domains are in similar positions relative to the saposin and that the enzyme active sites are adjacent to the saposin, although not directly overlaid (Fig. S4B). However, there are significant differences between these structures and this may reflect the different mechanisms by which the saposin domains facilitate lipid processing. Specifically, SapA is thought to function as a ‘solubiliser’, an assertion strongly supported by the GALC-SapA structure presented here. ASM, on the other hand, is likely to function via a ‘liftase’ mechanism. Specifically, ASM has been shown to bind tightly to negatively charged membranes and the saposin domain is likely to play a critical role in orienting the active site towards the membrane (33, 35).

Dimerisation of the saposin-like domain would not be required in this ‘liftase’ model of ASM activity, and indeed we observe that steric hindrance would prevent formation of a SapA-like dimer by the ASM saposin-like domain. We therefore conclude that the mechanisms used by saposins to facilitate lipid presentation will influence the stoichiometry and topology of their association with their cognate hydrolases.

The structure of GALC-SapA presented here defines how saposins can deliver sphingolipids to soluble hydrolases, presenting substrates to the hydrolase via a channel that shields the hydrophobic lipid tails from the surrounding polar solvent.

## Materials and Methods

### Protein expression and purification

His_6_-tagged murine GALC (mGALC) and untagged murine SapA (mSapA) were expressed and purified as described previously (28, 31). mSapA mutants were made by site-directed mutagenesis and verified by sequencing. Mutant proteins were expressed and purified exactly as for wild-type. Purified mGALC was concentrated to 5–15 mg/ml and stored in phosphate-buffered saline pH 7.4 at 4°C. Purified mSapA was concentrated to 8.0 mg/ml and stored in 50 mM Tris pH 7.4, 150 mM NaCl at 4°C.

### Pulldowns

mGALC was mixed with magnetic Ni-NTA agarose beads under saturating conditions (30 μl beads, 75 μg GALC per experiment) and incubated with mixing (90 min, 4 °C). Loaded beads were transferred to a flat bottomed 96-well plate and washed twice in pulldown buffer: 200 μl 100 mM sodium acetate pH 5.4, 20 mM NaCl, 0.05% Tween-20 or LDAO. For screening purposes the buffer pH, NaCl concentration and detergent were altered as appropriate. Concentrated SapA was pre-incubated with 0.1% Tween-20 or LDAO for 1 hour and 160 μg of SapA was added to GALC-loaded beads in 200 μl pulldown buffer and incubated with shaking (2 hr, 4 °C). Beads were then washed four times with 200 μl pulldown buffer. Proteins were eluted with 40 μl 500 mM imidazole, PBS pH 7.4, 0.05% Tween-20 and analysed by 4–12% gradient Bis-Tris PAGE. Following staining with Coomassie, gels were scanned on an Odyssey imaging system and band intensity determined using Image Studio Lite (LI-COR Biosciences).

### Protein complex crystallization

Purified mSapA was pre-incubated with LDAO in a final mix of 180 μM mSapA with 50mM sodium acetate pH 5.0, 700mM NaCl and 0.1% LDAO. This SapA-LDAO complex was incubated with purified GALC to give a final SapA:GALC molar ratio of 2:1 for crystallisation trials. Crystallization experiments were performed in 48-well sitting drops (800 nL of complex as prepared above plus 800 nL of precipitant) equilibrated at 20°C against 200 μL reservoirs of precipitant. Diffraction quality crystals grew against a reservoir of 75 mM sodium citrate pH 5.6 and 11% (w/v) PEG 3350. Crystals were cryoprotected in reservoir solution supplemented with 20% (v/v) glycerol and flash-cooled by plunging into liquid nitrogen.

### X-ray data collection, structure determination and refinement

Diffraction data were recorded at Diamond Light Source beamline I04-1 on a Pilatus 6M detector (Dectris). Diffraction data were collected at 100K. Data collection statistics are in Table 1. Diffraction data were indexed and integrated using DIALS and scaled and merged using AIMLESS *via* the xia2 automated data processing pipeline (36). Resolution cut-off was decided by CC_1/2_ value of >0.5 and *l*/ *σl* of 1.5 in the outer resolution shell.

The structure was solved by molecular replacement using Phaser with mouse GALC (PDB ID 4CCE (28)) and a model of mouse SapA, based on the structure of the human SapA in a lipo-protein disc (PDB ID 4DDJ (37)), as search models. Manual model building was performed using COOT (38). The structure was refined using autobuster (39), over-fitting of the data being minimised by the use of local structure similarity restraints (40) to the high-resolution structure of mouse GALC (PDB ID 4CCE (28)). Model geometry was evaluated with MolProbity throughout the refinement process (41). Final refinement statistics are presented in Table 1. Structural figures were rendered using PyMOL (Schrödinger LLC).

### Structure Analysis

The analysis of buried surface area and interface interactions in the GALC-SapA structure was carried out using the ePISA service at the European Bioinformatics Institute, EBI (42). Hydrogen bonding representations were created using LIGPLOT+ implementing DIMPLOT to generate schematic diagrams of protein-protein interactions (43). Structure-based alignments were carried out using SSM superposition implemented within COOT (44). Multiple sequence alignments were carried out using the MUSCLE service at the EBI (45). The atomic coordinates and structure factors for the GALC-SapA complex have been deposited in the Protein Data Bank (PDB ID 5N8K).

## Acknowledgements

We acknowledge Diamond Light Source for time on beamlines I04 and I04-1 under proposal MX11235. Remote access was supported in part by the EU FP7 infrastructure grant BIOSTRUCT-X (contract no. 283570). The Cambridge Institute for Medical Research is supported by Wellcome Trust Strategic Award 100140. J.E.D. is supported by a Royal Society University Research Fellowship (UF100371). This work is supported by MRC grant MR/N020626/1. S.C.G. is supported by a Sir Henry Dale Fellowship from the Royal Society and Wellcome Trust (098406/Z/12/Z).

